# A planarian nidovirus expands the limits of RNA genome size

**DOI:** 10.1101/299776

**Authors:** Amir Saberi, Anastasia A. Gulyaeva, John L. Brubacher, Phillip A. Newmark, Alexander E. Gorbalenya

## Abstract

RNA viruses are the only known RNA-protein (RNP) entities capable of autonomous replication (albeit within a permissive environment). A 33.5-kb nidovirus has been considered close to the upper size limit for such entities; conversely, the minimal cellular DNA genome is ~200 kb. This large difference presents a daunting gap for the transition from primordial RNP to contemporary DNA-RNP-based life. Whether or not RNA viruses represent transitional steps on the road to DNA-based life, studies of larger RNA viruses advance our understanding of size constraints on RNP entities. For example, emergence of the largest previously known RNA genomes (20-34 kb in positive-stranded nidoviruses, including coronaviruses) is associated with a proofreading exoribonuclease encoded in the nidoviral open reading frame 1b (ORF1b). However, apparent constraints on the size of ORF1b, which encodes this and other key replicative enzymes, have been hypothesized to limit further expansion of viral RNA genomes. Here, we characterize a novel nidovirus (planarian secretory cell nidovirus; PSCNV) whose disproportionately large ORF1b-like region, and overall 41.1 kb genome, substantially extend the presumed limits on RNA genome size. This genome encodes a predicted 13,556-aa polyprotein in an unconventional single ORF, yet retains canonical nidoviral genome organization and expression, and key replicative domains. Our evolutionary analysis suggests that PSCNV diverged early from multi-ORF nidoviruses, and subsequently acquired additional genes, including those typical of large DNA viruses or hosts. PSCNV’s greatly expanded genome, proteomic complexity, and unique features – impressive in themselves – attest to the likelihood of still-larger RNA genomes awaiting discovery.

**Significance Statement:** RNA viruses are the only known RNA-protein (RNP) entities capable of autonomous replication. The upper genome size for such entities was assumed to be <35 kb; conversely, the minimal cellular DNA genome is ~200 kb. This large difference presents a daunting gap for the proposed evolution of contemporary DNA-RNP-based life from primordial RNP entities. Here, we describe a nidovirus from planarians, whose 41.1 kb genome is 23% larger than the largest known of RNA virus. The planarian secretory cell nidovirus has broken apparent constraints on the size of the genomic subregion that encodes core replication machinery, and has acquired genes not previously observed in RNA viruses. This virus challenges and advances our understanding of the limits to RNA genome size.

## Introduction

Radiation of primitive life over its initial two billion years of evolution was likely accompanied by genome expansion and a proposed transition from RNA-based through RNA-protein to DNA-based life (1). The feasibility of an autonomous ancient RNA genome and the mechanisms underlying such fateful transitions are challenging to reconstruct. It is especially unclear whether such RNA entities ever evolved genomes close to the 100–300 kilobases (kb) range (2, 3) of the “minimal” reconstructed cellular DNA genome (4). This range overlaps with the upper size limit of nuclear pre-mRNAs (5); however, pre-mRNAs are incapable of self-replication, the defining property of primordial genomic RNAs.

RNA viruses may uniquely illuminate the evolutionary constraints on RNA genome size, whether or not they descended directly from primitive RNA-based entities (6–8). The average-sized RNA virus genome is around 10 kb, due to low-fidelity RNA replication without proofreading (9) However, positive-stranded RNA genomes of viruses of the order *Nidovirales*, which includes pathogenic coronaviruses, range from 12.7 to 33.5 kb, the largest known RNA genome (10, 11) (Fig. 1A, Table S1). These viruses may have evolved genomes larger than 20 kb after acquiring a proofreading exoribonuclease (ExoN) that improved replication fidelity (12–17). Despite this notable innovation, over the last 20 years of virus discovery, including unbiased metagenomics of invertebrate and vertebrate RNA viruses (18, 19), the largest-known RNA viral genome has only increased ~10% in size – a mere fraction of the nearly ten-fold increase observed for DNA viruses (20–22) (Fig. 1A). Apparently, genome sizes of large nidoviruses are tightly constrained by the interplay of genetic and host-specific factors, with the size of open reading frame 1b (ORF1b), encoding key replicative enzymes, emerging as a major barrier for further genome expansion (23). Hence, ~34 kb is considered the de facto limit of RNA virus genome size (18).

**Fig. 1.**
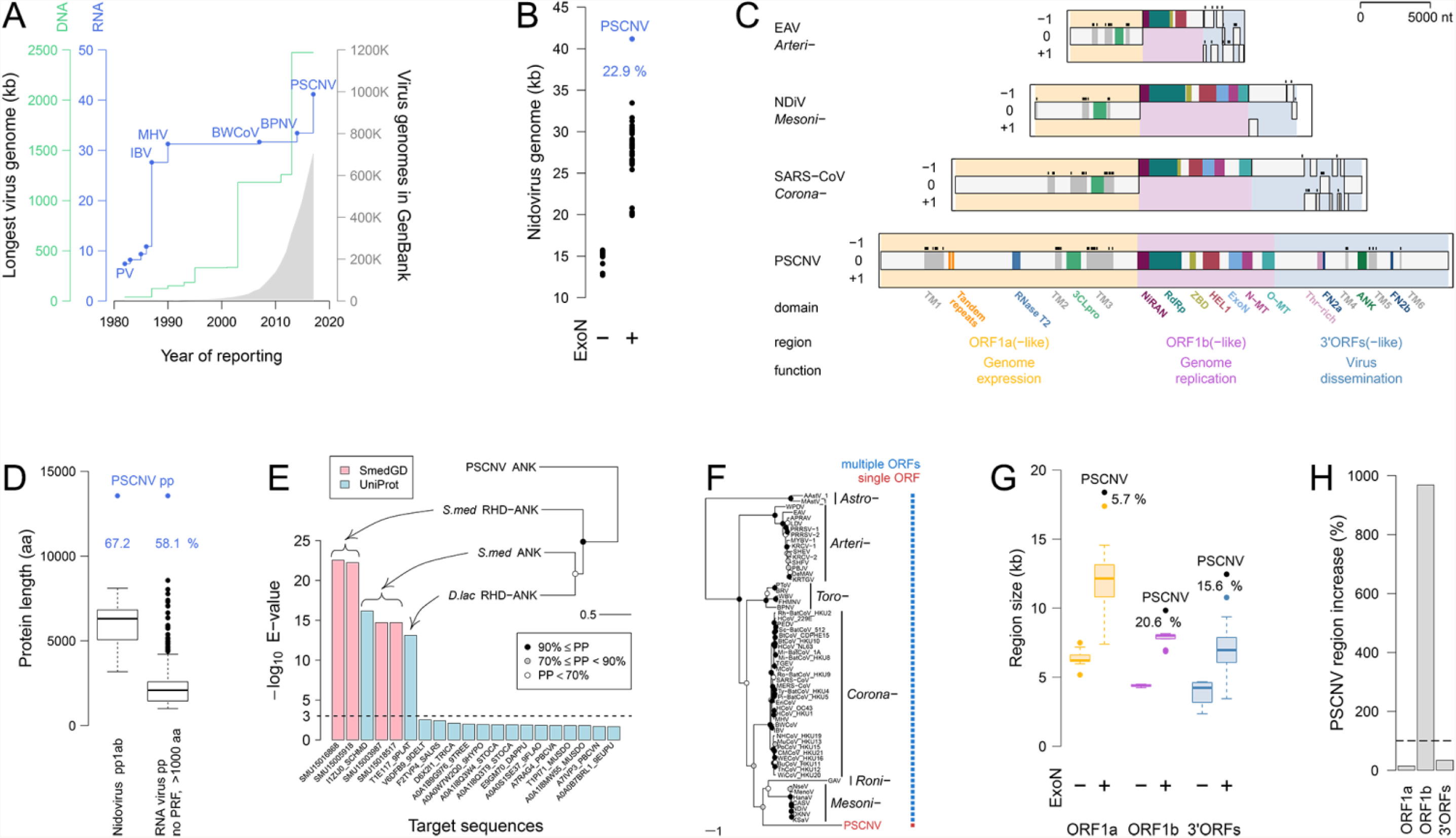
Genome and proteome of PSCNV. (***A***) Timeline of discovery of largest RNA and DNA virus genomes (1982–2017). PV, poliovirus. (***B***) Genome sizes of nidoviruses encoding an ExoN domain or not. Percentage indicates the difference between sizes of PSCNV and the next-largest entity (also in *D*,*G*). (***C***) ORFs and conserved protein domains in PSCNV and nidoviruses. ORF1a frame is set as zero. TM, transmembrane domain (TM helices shown by black bars above TM domains); ANK, Ankyrin; RNase T2, ribonuclease T2 homolog; for other domain acronyms, see text. (***D***) PSCNV polyprotein vs RNA virus protein sizes. (***E***) Closest cellular homologs of PSCNV ANK are ranked by similarity (left, above the broken baseline) and depicted through phylogeny (right; PP, posterior probability of clades) along with protein domain architecture: *S. med*, *Schmidtea mediterranea*; *D. lac*, *Dendrocoelum lacteum*; RHD, Rel homology DNA-binding domain. (***F***) Phylogeny and ORF number of PSCNV, nido-, and astroviruses (outgroup). (***G***) Sizes of three nidovirus ORF regions. (***H***) Size increase of three genome regions in PSCNV (grey bars) relative to the increase expected if all regions expanded evenly (broken line).

## Results and discussion

### Identification of a large RNA virus from planarians

To examine whether this limit applies beyond the currently recognized ~3000 RNA virus species that infect just hundreds of host species, further sampling of virus diversity is required. To this end, we analyzed *de novo* transcriptomes of the planarian *Schmidtea mediterranea* (24), in which viruses remain virtually unknown, and which exists as two reproductively distinct strains: sexuals that reproduce as simultaneous hermaphrodites and asexuals that reproduce via transverse fission (25). We determined the complete genome sequence of a putative nidovirus corresponding to one particularly large contig (SI Text, Table S2, Fig. S1). Excluding the polyadenylated tail, the fully assembled genome contains 41,103 nucleotides (nt) and encodes a single 40,671-nt ORF that is flanked by a 128-nt 5’-UTR and a 304-nt 3’-UTR (Fig. 1B,C). The presence of this RNA in vivo was confirmed by RT-PCR amplification of overlapping regions of 4–7 kb, 5’-RACE, and multiple EST clones (Fig. S1) (26). These sequences could not be amplified from *S. mediterranea* genomic DNA, nor could they be found in the genome (27); thus, they appear to derive from an exogenous source. Based on several criteria (see below), we assigned this RNA sequence to the genome of a virus we dubbed Planarian Secretory Cell Nidovirus (PSCNV). PSCNV circulates in both sexual and asexual host strains: analysis of RNA-seq data from three laboratories revealed its very low (<0.1%), albeit predominantly non-synonymous variation (SI Text, Tables S3,4). Analysis of *Planaria torva* transcriptome identified three contigs of the cumulative ~9400 nt size, whose regions displayed from 25% to 48% aa identity with PSCNV, indicative of at least one additional putative nidovirus (SI Text, Fig. S2) (28).

### The PSCNV proteome reveals a unique nidovirus

The genome and proteome of PSCNV are by far the largest yet reported for an RNA virus. Its RNA genome is ~25% larger than that of the next-largest known RNA virus (also a nidovirus), which is separated by a comparable margin from the first nidovirus genome sequenced 30 years ago (Fig. 1A). The size of the predicted PSCNV polyprotein (13,556 amino acids, aa) is 58–67% larger than the largest known RNA virus proteins produced from a single ORF (8,572 aa) or multiple ORFs through frameshifting (8,108 aa) (Fig. 1D).

We delineated at least twenty domains in the PSCNV polyprotein using a complex multistage procedure that combined different analyses within a probabilistic framework (Fig. 1C; SI Text, Table S5, and Figs. S3–S16). These domains include distant homologs of all ten domains characteristic of invertebrate nidoviruses (15, 29) and established with utmost confidence: seven ORF1a/1b enzymes (3C-like protease, 3CLpro; nidovirus RdRp-associated nucleotidyltransferase, NiRAN; RNA-dependent RNA polymerase, RdRp; superfamily 1 helicase, HEL1; exoribonuclease of DEDDh subfamily, ExoN; and SAM-dependent N7- and 2’-O-methyltransferases, N-MT and O-MT, respectively) and three non-enzymatic domains (transmembrane domain 2 and 3, TM2 and TM3, respectively; and zinc-binding domain, ZBD) (Fig. 1C). These domains are syntenic in PSCNV and nidoviruses, with the conservation of NiRAN and ZBD in PSCNV being of particular significance, since both domains are the only known genetic markers of the order *Nidovirales* (29, 30). Thus, these findings further support the phylogenetic affinity of PSCNV and nidoviruses (Fig. 1C), which we quantified in separate analyses (see below). Accordingly, three functional regions corresponding to the ORF1a, ORF1b, and 3’ORFs of nidoviruses could be identified tentatively in the PSCNV genome, with three characteristic nidovirus domains mapping to the ORF1a-like region and seven domains to the ORF1b-like region.

The enzymatic domains contain most essential catalytic residues, suggesting that they retain canonical functions, although notable deviations were also observed (Figs. S7–S14). Unique features of PSCNV’s putative enzymes include: a) 3CLpro encodes a valine (Val) residue in the position commonly occupied by a His residue in the putative substrate-binding pocket (GXV vs G/YXH) that controls the P1 substrate preference for Glu or glutamine (Gln) residues (31–35); b) a Ser residue of the nidovirus-specific SDD signature of RdRp (12) is replaced by glycine (Gly) – a characteristic of ssRNA+ viruses other than nidoviruses; c) the invariant aspartate (Asp), along with the highly conserved Ser/Thr, of motif B_N_ of the NiRAN (29) are substituted with Thr and Asn, respectively; d) four Cys/His residues coordinating a Zn^2+^ in the active site of ExoN (16, 17) are replaced in PSCNV, which, like non-nidoviral homologs, apparently lacks this Zn-finger.

Conserved sequence features reveal that, as in other nidoviruses (12, 36), the PSCNV 3’ORFs-like region may encode major structural proteins of enveloped virions, including the transmembrane glycoproteins known as S and M in coronaviruses, and nucleocapsid (N) protein (SI Text, Fig. S4). This region also appears to encode several notable domains: a ~130 aa region with Thr and Ser accounting for 70.7% of residues (Fig. S4); an Ankyrin domain (Figs. 1E and S16) and two fibronectin type II domains (FN2a and FN2b; Fig. S15), unique for RNA and all viruses, respectively. The Thr-rich and Thr/Ser-rich regions have been implicated in mediating adherence of fungal and bacterial extracellular (glyco) proteins to various substrates (37, 38), suggesting a similar role in PSCNV. The Ankyrin domain might counteract host immune responses by suppressing the NF-kB signaling pathway, as observed in poxviruses (39), while FN2a/b could facilitate virus dissemination (SI Text).

Computational analysis suggests that PSCNV might have acquired the Ankyrin domain from a planarian host relatively late in evolution (Fig. 1E), while the origins of the FN2 domains remain uncertain. We also identified a ribonuclease T2 homolog upstream of the putative 3CLpro in the ORF1a-like region (Fig. 1C), in which all active-site residues are conserved (Fig. S6). Pestivirus and polydnavirus ribonuclease T2 homologs modulate cell toxicity and immunity (40, 41).

### PSCNV likely derived from multi-ORF nidoviruses and expanded disproportionately in the ORF1b-like region

To learn about the evolution of PSCNV, we inferred the rooted phylogeny of PSCNV based on the RdRp protein domain of nidoviruses and two astroviruses (Fig. 1F; SI Text). PSCNV predominantly clustered with other invertebrate nidoviruses in the Bayesian sample, basal to either mesoni- and roniviruses (54.7% of the trees), roniviruses (20.6%), or mesoniviruses (13.4%). Ancestral state reconstruction analysis indicated that the most recent common ancestor of nidoviruses contained multiple ORFs (Log Bayes Factor 6.06) (Fig. S17). Thus, the single-ORF organization of PSCNV likely emerged through loss of signals separating the three ORF regions of ancestral nidoviruses.

Compared to known nidoviruses, PSCNV genome size increased unevenly in the three main regions (Fig. 1G,H, SI Text, Table S6). The size increase of the ORF1b-like region was an extraordinary 9-fold higher than would be expected from the extremely low variation in the size of this region among other nidoviruses normalized to genome size variation. This observation implies that PSCNV broke the tightly constrained ORF1b size barrier (23), but expanded modestly in the ORF1a-like and 3’ORFs-like regions. This pattern fits the wavelike dynamic model of genome expansion in nidoviruses (23). Accordingly, extant nidovirus genomes of different sizes reached particular points on a common trajectory of genome expansion dominated by consecutive increases of ORF1b, ORF1a, and 3’ORFs. A complete cycle of genome expansion was reported to encompass the 12.7-31.7 kb range, with the expansion of ORF1b expected to lead a second cycle beyond that size range. The sizes of PSCNV’s three main regions support this model, and show that nidoviruses were able to start a second cycle of genome expansion, whose feasibility has remained uncertain so far. This observation makes plausible the existence of other yet-undiscovered nidoviruses that have moved further along this genome-expanding trajectory.

### PSCNV infects the secretory cells of planarians

We next examined PSCNV infection in planarian tissues by whole-mount in situ hybridization (ISH). PSCNV RNA was detected abundantly in cells of the secretory system in both sexuals and asexuals (Fig. 2A). Fluorescent ISH revealed viral RNA in gland cell projections that form secretory canals (Fig. 2B). Notably, viral RNA was detected largely in ventral cells (Fig. 2C) whose localization corresponds to mucus-secreting cells producing the slime upon which planarians glide, and use to immobilize prey (42).

**Fig. 2.**
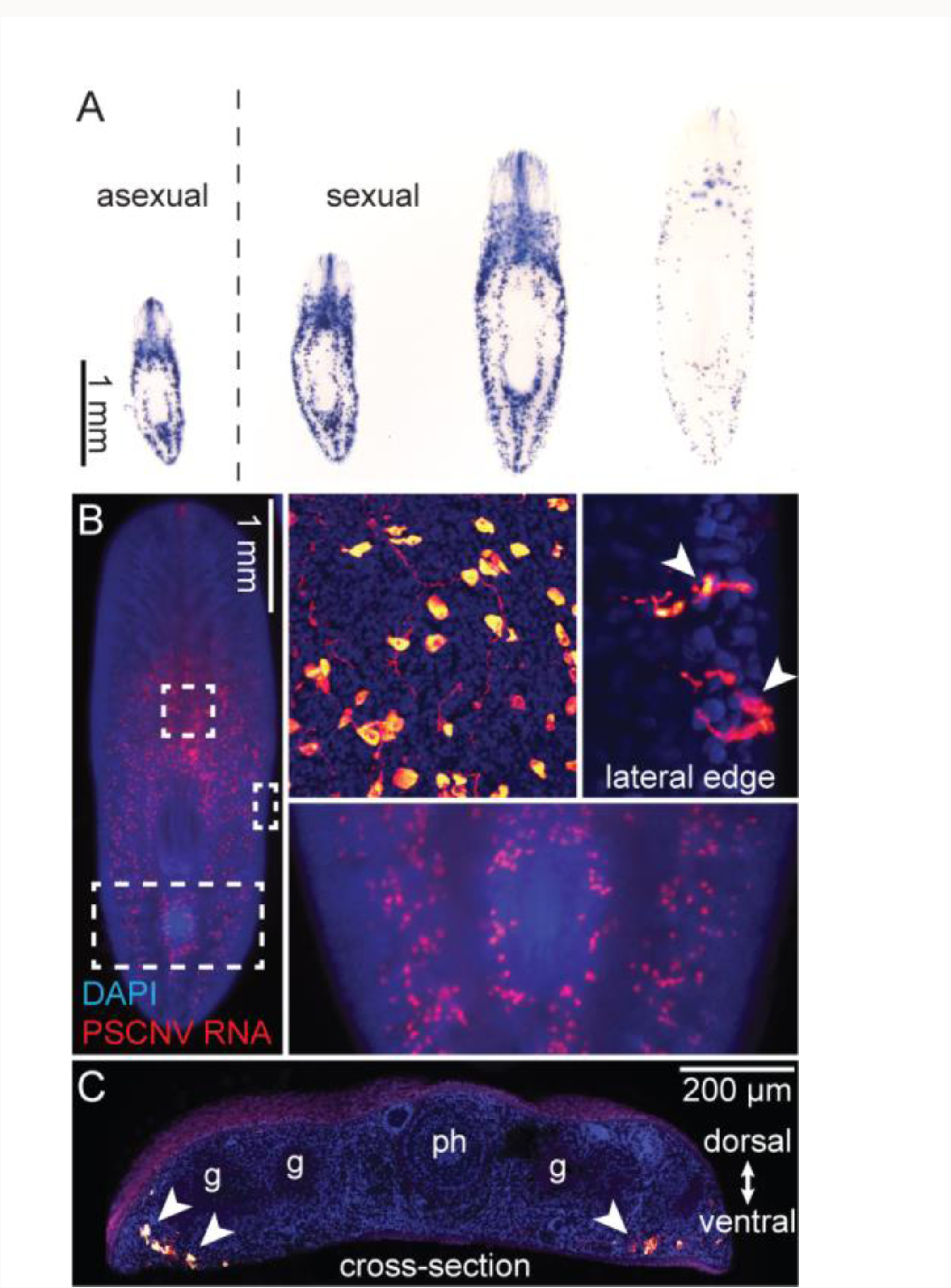
Expression of PSCNV RNA in planarians. (***A***) PSCNV RNA (blue) detected in asexual (left) and sexual *S. mediterranea* by whole-mount ISH. (***B***) Fluorescent ISH showing PSCNV expression in a sexual planarian. Insets show higher magnification of areas indicated by boxes. Top two insets are confocal projections. Secretory cell projections to lateral body edges are indicated by arrowheads. (***C***) Tiled confocal projections of PSCNV expression in a cross-section. Cells expressing PSCNV are ventrally located (arrowheads). Gut (“g”) and pharynx (“ph”) are indicated. DAPI (blue) labels nuclei.

We then analyzed planarians by electron microscopy (EM) for the presence of viral structures. Membrane-bound compartments with 90–150 nm spherical-to-oblong particles resembling nidoviral nucleocapsids (43, 44) were found in the cytoplasm of mucus-secreting cells, which are abundant in planarian parenchyma. These sub-epidermal gland cells are notable for their extensive rough endoplasmic reticulum and long projections into the ventral epithelium, through which they secrete mucus (Fig. S18). These cells provide an ideal environment for nidoviral replication – which co-opts host membranes to produce viral replication complexes (45). Putative viral particles were found both in deep regions of these cells, and in their trans-epidermal projections (Fig. 3A–C). The latter location suggests a route for viral transmission. Notably, particles in sub-epidermal layers have a “hazy” appearance and are embedded in a relatively electron-dense matrix (Fig. 3D). In contrast, particles closer to the apical surface of the epidermis appear as relatively discrete structures, standing out against electron-lucent surrounding material (Fig. 3E). The size, ultrastructure, and host-cell locations are all consistent with these structures being nidoviral nucleocapsids (43, 44).

**Fig. 3.**
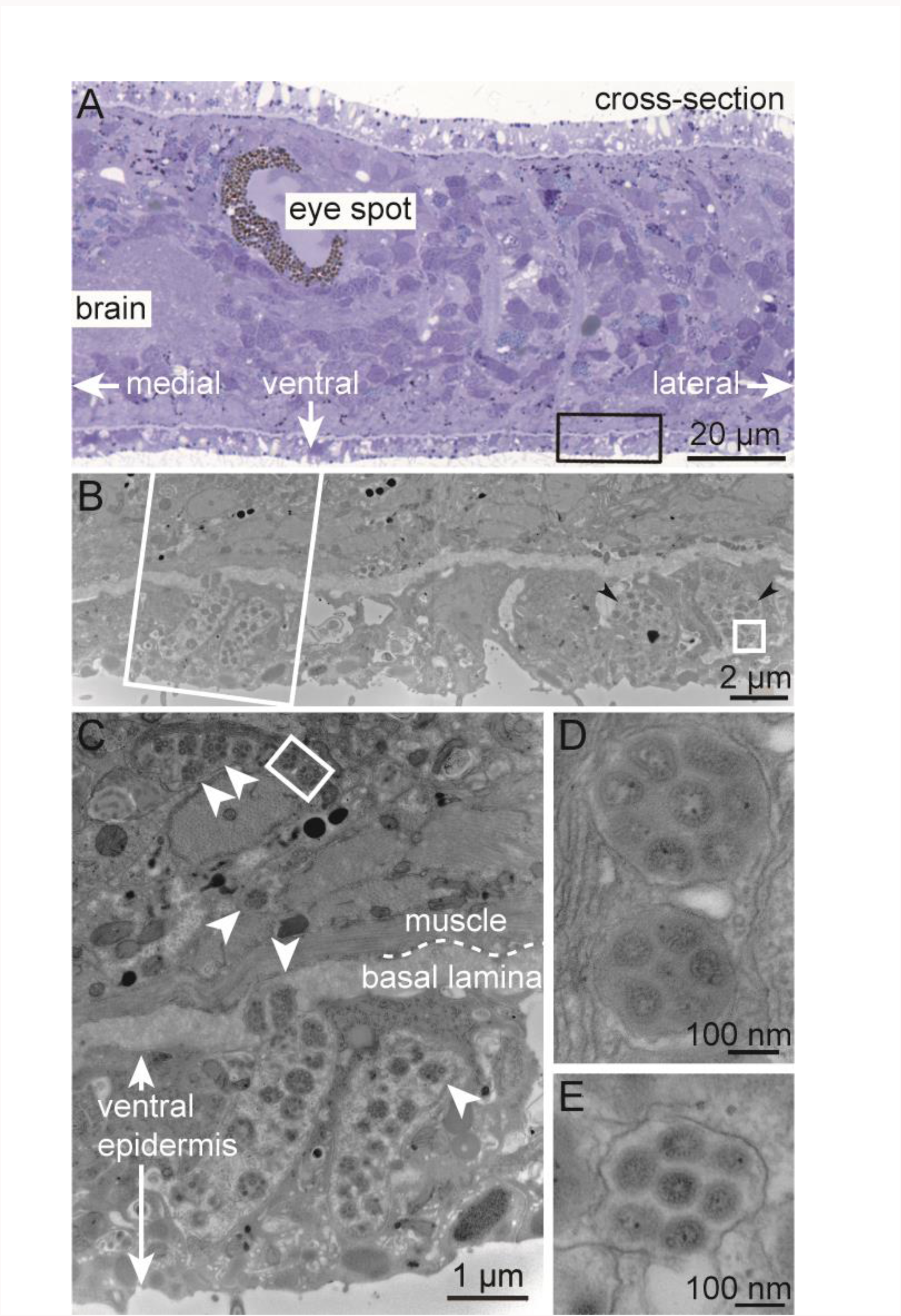
Putative PSCNV particles revealed by electron microscopy. (***A***) Adjacent histological section to orient EM images. Black rectangle corresponds to location of (***B***), a low-magnification EM view to provide context. White rectangle corresponds to location of (***C***), in which putative viral particles enclosed within membrane sacs are indicated by arrowheads. White rectangle in (*C*) and square in (*B*) indicate positions of higher magnification views shown in (***D***) and (***E***), respectively – each illustrating several viral particles within a membrane sac. In top-left of (*B*), note the mucus granules adjacent to virus-laden sacs (see also Fig. S18). Scale bars as indicated.

### Potential mechanisms for varying stoichiometries of viral proteins encoded by a single-ORF genome

Virus reproduction requires different viral protein stoichiometries at distinct replicative cycle stages, a challenge for a single-ORF genome theoretically producing equimolar quantities of encoded polypeptides. For example, the putative structural proteins encoded by 3’ORFs are needed in much larger quantities than replicative enzymes at the end of nidovirus infection, and vice versa. We analysed the PSCNV genome and its transcript expression to resolve this apparent paradox.

Nidoviruses use −1 programmed ribosomal frameshifting (PRF) at the ORF1a/1b border to attenuate the synthesis of major replicative enzymes upon genome translation (46). We located a potential −1 PRF signal in the canonical position in the PSCNV genome (SI Text). This signal could potentially attenuate in-frame translation downstream of the ORF1a-like region (Fig. 4A) similar to a mechanism used by encephalomyocarditis virus (47).

**Fig. 4.**
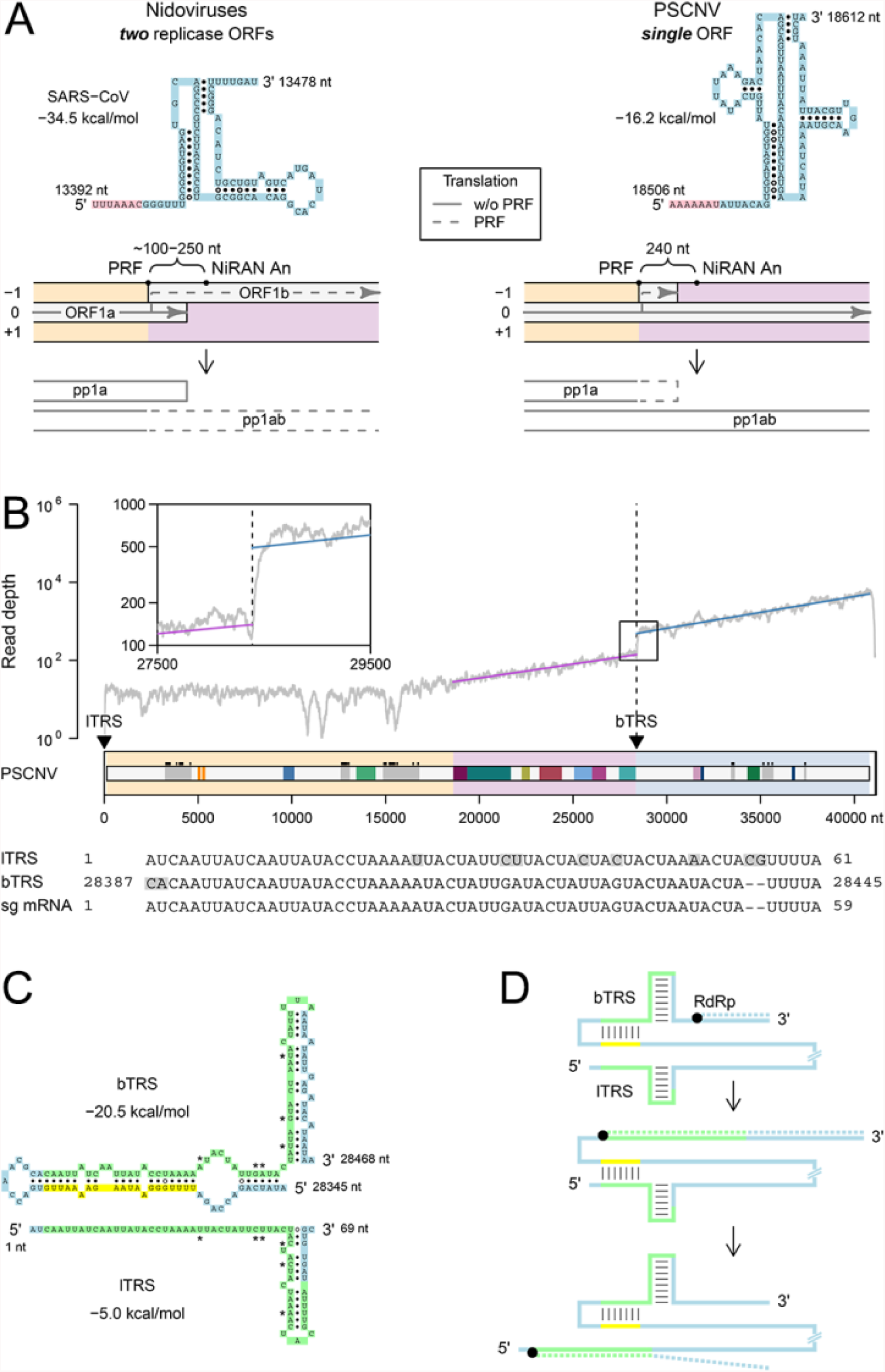
Genome translation and transcription. (***A***) Comparison of replicase ORFs translation mechanism in nidoviruses (left) and PSCNV (right, hypothetical). On the top, RNA structure of the PRF sites is presented: slippery sequence, pink; pseudoknot, blue. (***B***) Mean depth of RNA-seq coverage along the PSCNV genome (approximated by exponential regression in ORF1b-like and 3’ORFs-like regions) calculated based on five datasets used to assemble the transcriptomes in which PSCNV was found (24), position of oligonucleotide repeats (leader and body TRSs) in the genome, and their alignment with subgenomic (sg) mRNA 5’-terminus established by 5’-RACE (nucleotide mismatches between sg mRNA and TRSs are shown on grey background). (***C***) Predicted secondary structure of TRSs. TRSs are highlighted in green, region upstream of bTRS, interacting with its 5’-terminus in yellow, asterisks indicate mismatching nucleotides of TRSs. (***D***) Model of discontinuous RNA synthesis mediated by TRSs and their secondary structure. Genome is shown by solid line and nascent minus strand by dashed line. Color code matches that of panel *C*.

The PSCNV putative 3CLpro domain is expected (31) to release itself and all downstream polyprotein components up to the border between the ORF1b- and 3’ORFs-like regions (Fig. 1C). The polyprotein region processed by 3CLpro, and its flanking regions, are of comparable sizes (5006 vs 4400 and 4150 aa), suggesting further posttranslational processing. Indeed, we have identified a number of candidate cleavage sites for host furin-like proteases (Fig. S4), commonly involved in processing viral glycoprotein precursors (48) (SI Text).

We obtained several lines of evidence for upregulated transcription of the 3’ORFs-like region (Fig. 4B–D; SI Text). The PSCNV genome contains a pair of 59 and 57-nt imperfect repeats with 86% identity starting at positions 3 and 28,389, respectively (Fig. 4B); their positioning immediately upstream of ORF1a-like and at the border between ORF1b- and 3’ORFs-like regions resembles that of leader and body transcription-regulating sequences (l- and bTRSs) in other nidoviruses (49). Accordingly, we found a sharp (~3-fold) change in RNA-seq read coverage at the bTRS. We identified an RNA species (by 5’-RACE), whose first two and following 57 nucleotides matched the genomic 5’-terminal AU followed by the bTRS (Fig. 4B), suggesting the presence of a subgenomic mRNA produced by discontinuous transcription. Furthermore, the bTRS potentially forms strong complementary interactions with its upstream and downstream sequences (Fig. 4C). When unwound, the upstream sequence can form a duplex with the lTRS, bringing together the two spatially separated genomic regions, consistent with a model of discontinuous subgenomic RNA synthesis (Fig. 4D) resembling that of other nidoviruses (50).

## Conclusions

The advent of metagenomics and transcriptomics have greatly accelerated the pace of virus discovery, leading to studies reporting genome sequences of dozens to thousands of new RNA viruses in poorly characterized hosts (18, 19, 51, 52). These developments have substantially advanced our appreciation of RNA virus diversity, and improved our understanding of the mechanisms of its generation (53, 54). Notwithstanding that sea change, the largest known RNA genomes continue to belong to nidoviruses, as has been the case for 30 years since the first coronavirus genome of 27 kb was sequenced (11, 15, 55) (Fig. 1A). The unbiased, transcriptomics-based discovery of PSCNV in planarians reinforces the status of nidoviruses as relative giants among RNA viruses, while demonstrating that RNA genomes may be substantially larger than previously understood. Further, genomic and proteomic characteristics of PSCNV defy the perceived central role of multiple ORFs in the life cycle and evolution of nidoviruses. Contrary to conventional wisdom, single-ORF genome expression may involve subgenomic mRNA synthesis, while functional constraints linked to the synteny of key replicative enzymes may be the hallmark characteristics of nidoviruses. Since the ORF1b-size barrier can be overcome and genome organization changed, nidoviruses of yet-to-be-sampled hosts might prove to have evolved even larger RNA genomes than that reported here, further decreasing the gap between virus RNA and host DNA genome sizes.

## Materials and Methods

All Materials and Methods are described in Supplementary Information in detail.

### PSCNV genome and its variants in *S. mediterranea* RNA-seq data

PSCNV genome was assembled based on contigs identified in two *de novo* transcriptomes of *S. mediterranea* (24) by comparison with genome of human coronavirus OC43 (KY014282.1) using tblastx (BLAST+ v2.2.29 (56)); termini of the assembly were completed by contigs from the two transcriptomes and planarian EST clones (26), identified by similarity to the incomplete assembly using blastn, and 5’-RACE amplicons. Reads from planarian RNA-seq datasets: in-house ones used to assemble the two transcriptomes, and the ones available from EBI ENA (57), were mapped to the PSCNV genome sequence by either CLC Genomics Workbench 7, or Bowtie2 version 2.1.0 (58); read counts and coverage were estimated using SAMtools 0.1.19 (59), genome sequence variants were called by BCFtools 1.4 (60).

### Reverse transcription, PCR, and 5’-RACE

Freshly prepared RNA from mature sexual planarians was used for cDNA synthesis (iScript, Bio-Rad) or 5’-RACE (RLM-RACE, Ambion) according to manufacturer instructions. Large overlapping amplicons across the PSCNV genome (primers in Table S2) were amplified by standard Phusion^®^ High-Fidelity DNA polymerase reaction with 65ºC primer annealing temperature and 10 min extension steps.

### In situ hybridization

Colorimetric and fluorescent in situ hybridizations were done following published methods (61). Digoxigenin (DIG)-labelled PSCNV probes were generated by antisense transcription of the planarian EST clone PL06016B2F06 (GenBank DN313906.1) (26). Following color development, all samples were cleared in 80% (v/v) glycerol and imaged on a Leica M205A microscope (colorimetric) or a Carl Zeiss LSM710 confocal microscope (fluorescent).

### Histology and Transmission Electron Microscopy

Sexual and asexual planarians originating from the Newmark laboratory were fixed and processed for epoxy (Epon-Araldite) embedding as previously described (62). For light-microscopic histology, 0.5 µm sections were stained with 1% (w/v) toluidine blue O in 1% (w/v) borax for 30 s at 100°C, and imaged on a Zeiss Axio Observer. For transmission electron microscopy, 50–70 nm sections were collected on copper grids, stained with lead citrate (63) and imaged with a AMT 1600 M CCD camera on a Hitachi H-7000 STEM at 75 kV.

### Genome and Protein databases

For various analyses we used the following databases: PlanMine (28), Smed Unigene (64), scop70_1.75, pdb70_06Sep14 and pfamA_28.0 supplemented with profiles of conserved nidovirus domains (65–67), Uniprot (68), list of genome sequences representing nidovirus species (69), NCBI Viral Genomes Resource (70), GenBank (71) and RefSeq (72).

### Computational protein and RNA sequence analyses

Virus protein sequences were analyzed to predict disordered regions (DisEMBL 1.5 (73)), transmembrane regions (TMHMM v.2.0), secondary structure (Jpred4 (74)), signal peptides (SignalP 4.1 (75)), N-glycosylation sites (NetNGlyc 1.0) and furin cleavage sites (ProP 1.0 (76)). To identify sites enriched with amino acid residue, distribution of each residue along polyprotein sequence was assessed using permutation test executed with a custom R script. Generation of multiple sequence alignments of RNA virus proteins was facilitated by Viralis platform (77). Protein homology profile-based analyses were assisted with HMMER 3.1 (78), and HH-suite 2.0.16 (79). To predict RNA secondary structure and PRF sites we used Mfold web server (80) and KnotInFrame (81), respectively. Blastn (BLAST+ v2.2.29) (56) was used to identify RNA repeats.

### Evolutionary analyses

Phylogeny was reconstructed by Bayesian approach using a set of tools including BEAST 1.8.2 package (82) and ProtTest 3.4 (83) as described in (84). BayesTraits V2 (85) was used to perform ancestral state reconstruction. Preference for a state at a node was considered statistically significant only if Log BF exceeded 2 (86).

### Visualization of results

Protein alignments were visualized with the help of ESPript 2.1 (87). To visualize Bayesian samples of trees, DensiTree.v2.2.1 was used (88). R was used for visualization (89).

## Data availability

Contigs used to assemble PSCNV genome were deposited to GenBank (accession nos. BK010447-BK010449).

## Acknowledgments

We thank Paul Ahlquist for helping initiate this collaboration and Johan den Boon, Andrey M. Leontovich, Dmitry V. Samborskiy, and Igor A. Sidorov for discussions and assistance. This work was supported by NIH R01 HD043403 to P.A.N. and by EU Horizon2020 EVAg 653316 project and LUMC MoBiLe program to A.E.G.; P.A.N. is an investigator of the Howard Hughes Medical Institute and A.E.G. was Leiden University Fund Professor at the time of this study.

